# TSPO deficiency reveals a novel role for porphyrins in regulating white adipose tissue lipid metabolism

**DOI:** 10.1101/2025.09.14.676078

**Authors:** Carmen R. Smith, Lan N. Tu, Prasanthi P. Koganti, Vimal Selvaraj

## Abstract

The translocator protein (TSPO) is a mitochondrial outer membrane protein with high expression in cells specialized for lipid metabolism, including white adipose tissue (WAT), where its physiological function remains poorly defined. Our previous work demonstrated a role for TSPO in mitochondrial fatty acid oxidation, prompting us to investigate its contribution to adipocyte lipid homeostasis under metabolic stress. We employed a combination of *in vivo* and *in vitro* approaches using global *Tspo* knockout (*Tspo^-/-^*) mice, primary adipocyte cultures, and pharmacological TSPO-binding drugs to examine TSPO function. We show that *Tspo^-/-^* mice exhibited increased WAT mass and adipocyte hypertrophy following high-fat diet feeding, with no changes in caloric intake. These changes were accompanied by suppression of key lipolytic genes, reduced circulating NEFA and glycerol levels, and altered expression of fatty acid oxidation genes. Lipidomic profiling showed no genotype-dependent changes, indicating impaired mobilization rather than altered lipid composition. *In vitro*, TSPO deficiency enhanced lipid accumulation during adipocyte differentiation and impaired expression of lipolytic genes. TSPO-binding drugs phenocopied this response in TSPO-expressing cells. Mechanistically, we identify protoporphyrin IX (PPIX), an endogenous TSPO ligand, as a suppressor of adipogenic and lipolytic programs. PPIX levels increase during adipogenesis, and its accrual inhibits both lipid accumulation and lipolytic response, whereas hemin does not elicit these effects. Our findings identify TSPO as a regulator of adipocyte lipid metabolism through a previously unrecognized TSPO-PPIX axis that influences lipid mobilization in adipocytes. This mechanism provides insight into TSPO’s role in metabolic adaptation and highlights its potential as a therapeutic target in obesity-associated adipose dysfunction.

## Introduction

The translocator protein (TSPO), formerly known as the peripheral benzodiazepine receptor (PBR), is an evolutionarily conserved 18-kDa protein embedded in the outer mitochondrial membrane (OMM) (Braestrup & Squires 1977; Anholt *et al*. 1986). Initially discovered in steroidogenic tissues based on its binding affinity for benzodiazepine-like ligands, TSPO was proposed to facilitate mitochondrial cholesterol import – a critical, rate-limiting step in steroid hormone biosynthesis (Mukhin *et al*. 1989; Papadopoulos *et al*. 1990). This model was supported by its OMM localization and the presence of a putative cholesterol recognition/interaction amino acid consensus (CRAC) motif (Anholt *et al*. 1986; Li *et al*. 2001). However, definitive genetic ablation studies have since unequivocally demonstrated that TSPO is dispensable for steroidogenesis in mammals (Banati *et al*. 2014; Morohaku *et al*. 2014; Tu *et al*. 2014), with mitochondrial cholesterol import now solely attributed to the intermembrane space shuttle function of the steroidogenic acute regulatory protein (STAR) (Koganti *et al*. 2025). Separately, TSPO was also found to associate with components of the mitochondrial permeability transition pore (MPTP), leading to a functional model linking TSPO to regulation of reactive oxygen species (ROS) production and apoptosis (McEnery *et al*. 1992); however, this model has similarly been disproven (Justina Šileikytė *et al*. 2014). These findings have necessitated a major reassessment of TSPO’s biological role, positioning it as a central subject of renewed investigation in mitochondrial physiology (Selvaraj & Stocco 2015; Gavish & Veenman 2018; Bréhat *et al*. 2024).

TSPO remains of considerable biomedical interest due to its altered expression across a wide range of pathological conditions, including cancer, neurodegenerative diseases, inflammation, and metabolic dysfunction (Galiègue *et al*. 2004; Vlodavsky & Soustiel 2007; Thompson *et al*. 2013; Jimenez *et al*. 2022). Notably, TSPO-targeted positron emission tomography (PET) imaging is widely employed to infer neuroinflammation (Mirzaei *et al*. 2016; Singh *et al*. 2022), further emphasizing its perceived involvement in cellular stress responses and mitochondrial signaling. Recent studies have proposed that TSPO may contribute to mitochondrial quality control, regulation of reactive oxygen species (ROS) production (Loth *et al*. 2020), apoptosis (Corsi *et al*. 2022), and cellular bioenergetics (Liu *et al*. 2017), although these models remain mechanistically unresolved and sometimes contradictory.

Previous work from our laboratory (Tu *et al*. 2016), and emerging transcriptomic and proteomic datasets (Uhlén *et al*. 2015) reveal that TSPO is highly expressed not only in classical lipid-rich steroidogenic organs/tissues (Koganti & Selvaraj 2020), but also in other metabolically active tissues, particularly white adipose tissue (Thompson *et al*. 2013). In rodents, adipose TSPO expression appears to be modulated by metabolic status, including diet-induced obesity, thermogenic stimulation, and nutrient availability (Hartimath *et al*. 2019). However, functional studies investigating TSPO’s contribution to lipid metabolism have been limited. Several reports suggest that TSPO expression may respond to metabolic stress and that its pharmacological ligands modulate lipid storage phenotypes (Gut *et al*. 2013; Kim *et al*. 2019), although the specificity and mechanism of these effects remain unclear. Given the pivotal role of mitochondria in adipocyte differentiation, lipid oxidation, and lipolysis, we considered the possibility that TSPO may directly modulate adipocyte lipid metabolism.

Unbiased discovery approaches have also implicated TSPO in adipose tissue lipid metabolism. In a genetic mapping study integrating quantitative trait loci (QTL) analysis with bioinformatic prioritization, TSPO was identified as a candidate gene influencing plasma triglyceride levels in mice (Leduc *et al*. 2011), suggesting that natural variation at the *Tspo* locus may affect systemic lipid homeostasis. Independently, an exploratory screen for adipogenesis-regulated genes using differential display RT-PCR also uncovered that expression of *Tspo* was markedly upregulated during the differentiation of 3T3-L1 preadipocytes into mature adipocytes (Wade *et al*. 2005). These early findings indicated that TSPO expression dynamically responds to adipogenic stimuli. Aligned with these findings, we previously demonstrated that TSPO regulates mitochondrial fatty acid oxidation in steroidogenic cells (Tu et al. 2016), linking TSPO activity to core aspects of mitochondrial lipid metabolism. Nevertheless, direct functional evidence for TSPO in white adipose tissue remains limited.

In this study, we leveraged a multifaceted experimental strategy involving global Tspo knockout (*Tspo^-/-^*) mice, primary adipocyte cultures, and pharmacological interventions to dissect the role of TSPO in white adipose tissue (WAT) function. We report that TSPO deficiency enhances lipid accumulation in adipocytes under metabolic stress, driven by suppression of canonical lipolytic pathways. Mechanistically, we identify the endogenous TSPO ligand PPIX as a potential mediator of this phenotype. Although TSPO is not essential for heme synthesis *per se* (Zhao *et al*. 2016), we show that PPIX concentrations rise during adipogenesis and that excess PPIX suppresses lipolysis and adipogenic gene expression. These data suggest a previously unrecognized role for TSPO in modulating adipocyte lipid metabolism via a PPIX-dependent mechanism, with potential implications for understanding obesity-associated adipose dysfunction and the development of TSPO-targeted therapies.

## Results

### TSPO deficiency increases white adipose tissue mass

To determine the impact of TSPO deficiency under conditions of metabolic stress, we compared body weight trajectories, adipose tissue mass, and feed intake in *Tspo^fl/fl^* and *Tspo^-/-^* mice subjected to high-fat (HF) diet feeding. At baseline (6 weeks of age), there were no significant differences in total body weight between genotypes in either males or females (Figure 1A). Over the 8-week HF diet period, male *Tspo^fl/fl^* and *Tspo^-/-^* mice maintained similar body weight trajectories, whereas female *Tspo^-/-^* mice exhibited a modest but significant increase in body weight relative to *Tspo^fl/fl^* female mice beginning at week 7 (Figure 1B). Despite these minor body weight differences, analysis of adipose tissue mass revealed a significant increase in the weight of gonadal (gWAT) and inguinal (iWAT) white adipose tissue depots in both male and female *Tspo^-/-^* mice compared to controls following HF diet feeding, while liver weights remained unchanged (Figure 1C). Daily feed intake during the HF diet period was comparable between genotypes (Figure 1D), indicating that the enhanced adipose tissue expansion observed in *Tspo^-/-^* mice was not attributable to differences in caloric intake.

**Figure 1.**
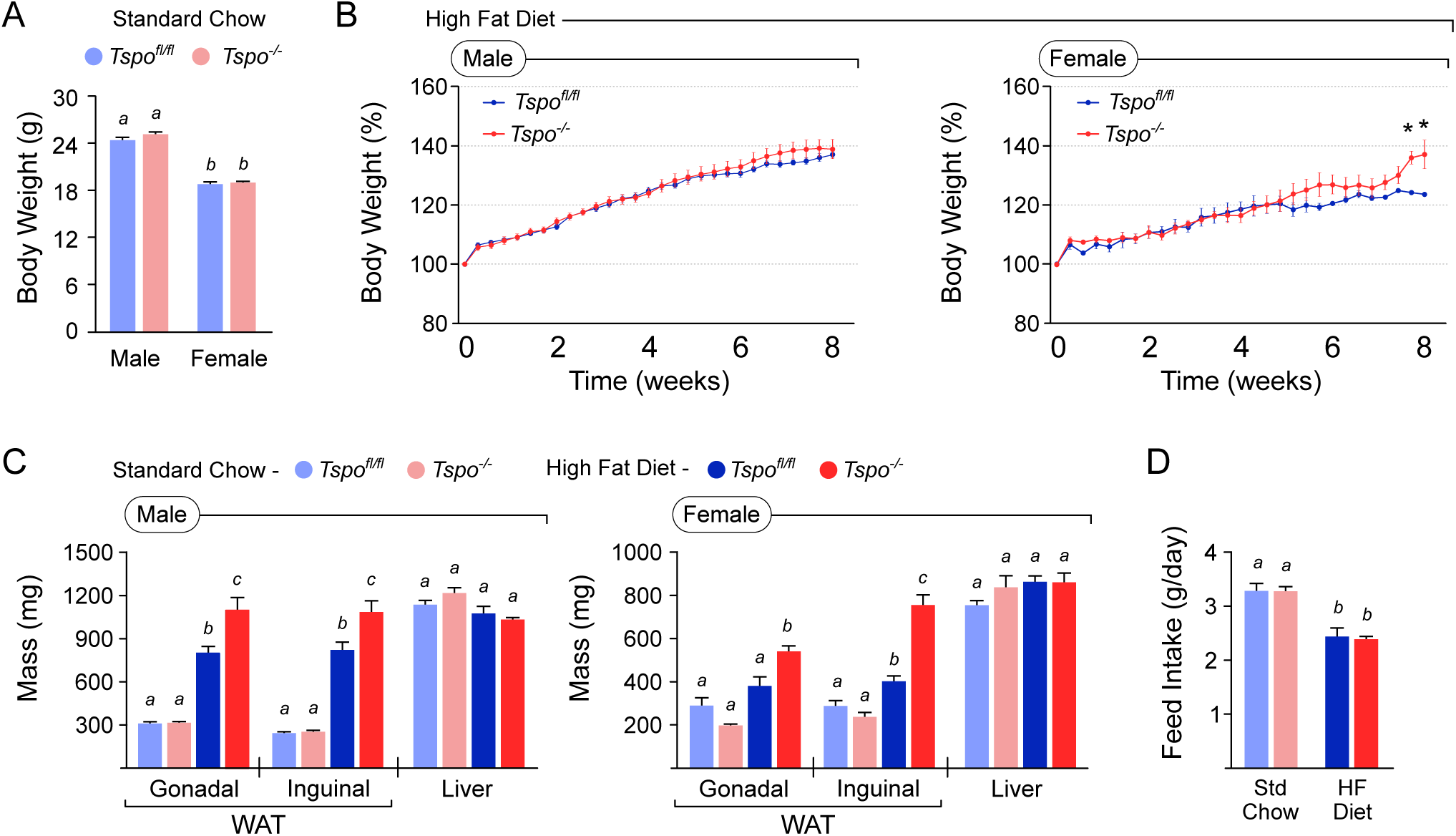
TSPO deficiency increases adipose depot mass without altering total body weight following high-fat diet feeding. (A) Body weights of *Tspo^fl/fl^* and *Tspo^-/-^* male and female mice at baseline showed no significant differences between genotypes (8 weeks old; n=10 per group). (B) Body weight gain trajectories over the 8-week high fat (HF) diet period revealed no significant genotype effect in males, but showed significant increases after 7 weeks in female *Tspo^-/-^* mice (*p<0.05; n=10 per group for males and females). (C) Quantification of white adipose tissue (WAT) depots indicated a significant increase in gonadal WAT (gWAT) and inguinal WAT (iWAT) weights in *Tspo^-/-^* mice compared to *Tspo^fl/fl^* controls in both sexes following HF diet feeding (n=10 per group), despite similar total body weights. Liver weights remained unchanged. (D) Daily feed intake (g/day) was not significantly different between *Tspo^fl/fl^* and *Tspo^-/-^* mice during the 8-week HF diet period, indicating that the enhanced WAT expansion was independent of differences in caloric intake. Different letters denote statistically significant differences (p < 0.05).

### TSPO deficiency promotes adipocyte hypertrophy

We next assessed whether TSPO deficiency alters adipocyte morphology under baseline and high-fat diet (HF) conditions. Histological analysis of gWAT sections from chow-fed mice revealed no overt differences in adipocyte size between *Tspo^fl/fl^* and *Tspo^-/-^* genotypes (Figure 2A). Quantification of adipocyte area and cell size distribution confirmed that TSPO deficiency did not significantly impact adipocyte size under standard chow conditions (Figure 2B). In contrast, after 8 weeks of HF diet feeding, gWAT sections from *Tspo^-/-^* mice showed visibly enlarged adipocytes compared to *Tspo^fl/fl^* controls (Figure 2C). Morphometric analysis demonstrated a significant increase in mean adipocyte area in *Tspo^-/-^* mice, accompanied by a rightward shift in the overall distribution of adipocyte sizes (Figure 2D). These findings indicate that loss of TSPO promotes adipocyte hypertrophy in response to high-fat diet-induced metabolic stress.

**Figure 2.**
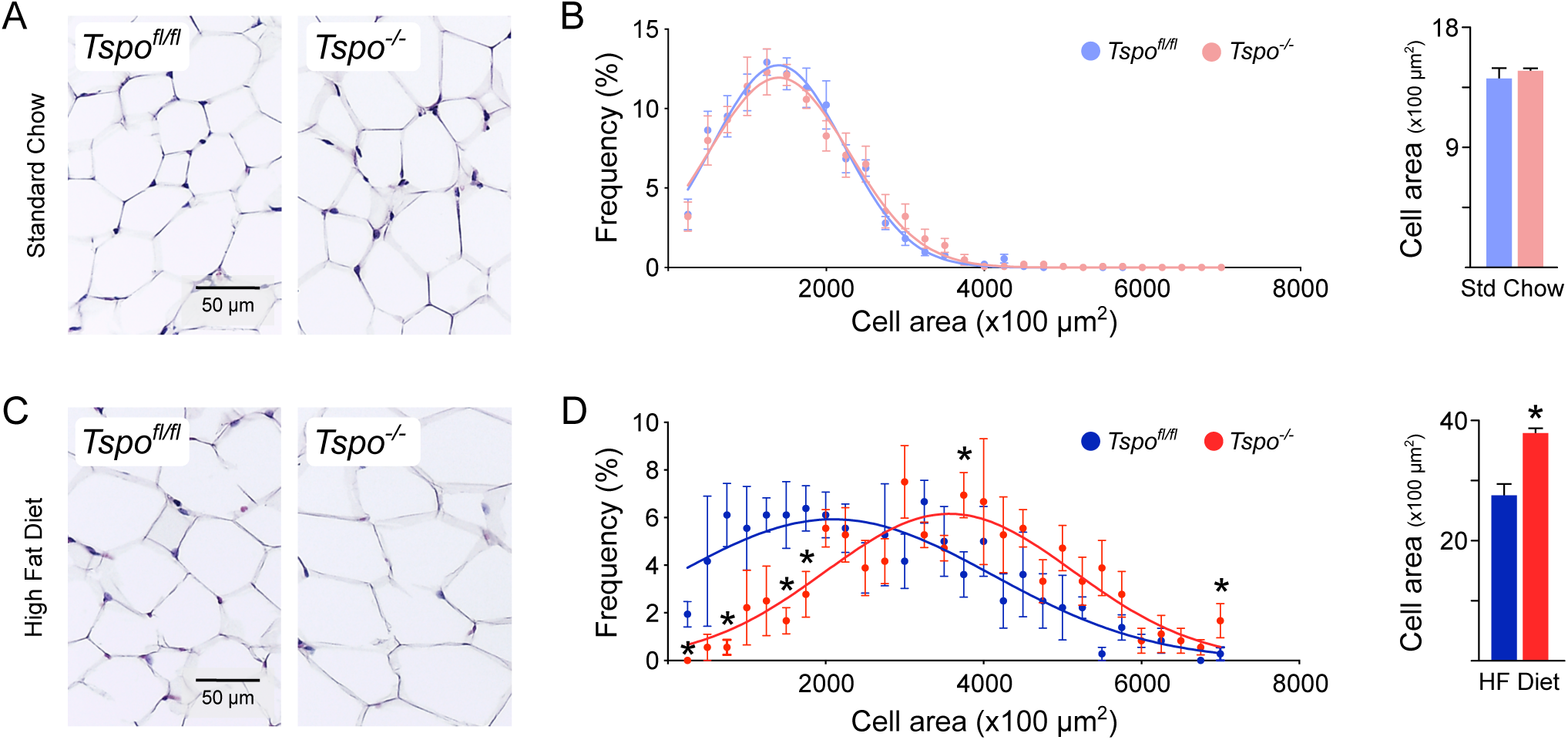
TSPO deficiency promotes adipocyte hypertrophy in white adipose tissue following high-fat diet feeding. (A) Representative hematoxylin and eosin-stained sections of gonadal white adipose tissue (WAT) from *Tspo^fl/fl^* and *Tspo^-/-^* mice under standard chow conditions. (B) Under standard chow, quantification of adipocyte area revealed no significant differences between genotypes, as shown by mean adipocyte area (right) and cell size distribution frequency (left; n=1344 cells per group). (C) Representative hematoxylin and eosin-stained sections of gonadal WAT from *Tspo^fl/fl^* and *Tspo^-/-^* mice following high-fat diet feeding shows adipocyte hypertrophy. (D) Following 8 weeks of HF diet, *Tspo^-/-^* adipocytes were significantly larger than *Tspo^fl/fl^* adipocytes, with increased mean cell area (left) and a rightward shift in cell size distribution (right; n=360 cells per group). Statistical significance for mean adipocyte area comparisons and distribution comparisons is indicated (*p < 0.05).

### Loss of TSPO alters lipid metabolism gene expression in WAT

To investigate the metabolic processes underlying the enhanced lipid accumulation observed in *Tspo^-/-^* mice, we analyzed the expression of genes involved in nutrient uptake, fatty acid (FA) synthesis, FA oxidation, lipolysis, and adipokine secretion in gWAT from mice fed either standard chow or HF diets. For genes involved in nutrient uptake, *Tspo^-/-^* WAT under chow diet did not exhibit significant changes in the expression of *Cd36*, *Lpl*, or *Glut4* compared to controls (Figure 3A).

**Figure 3.**
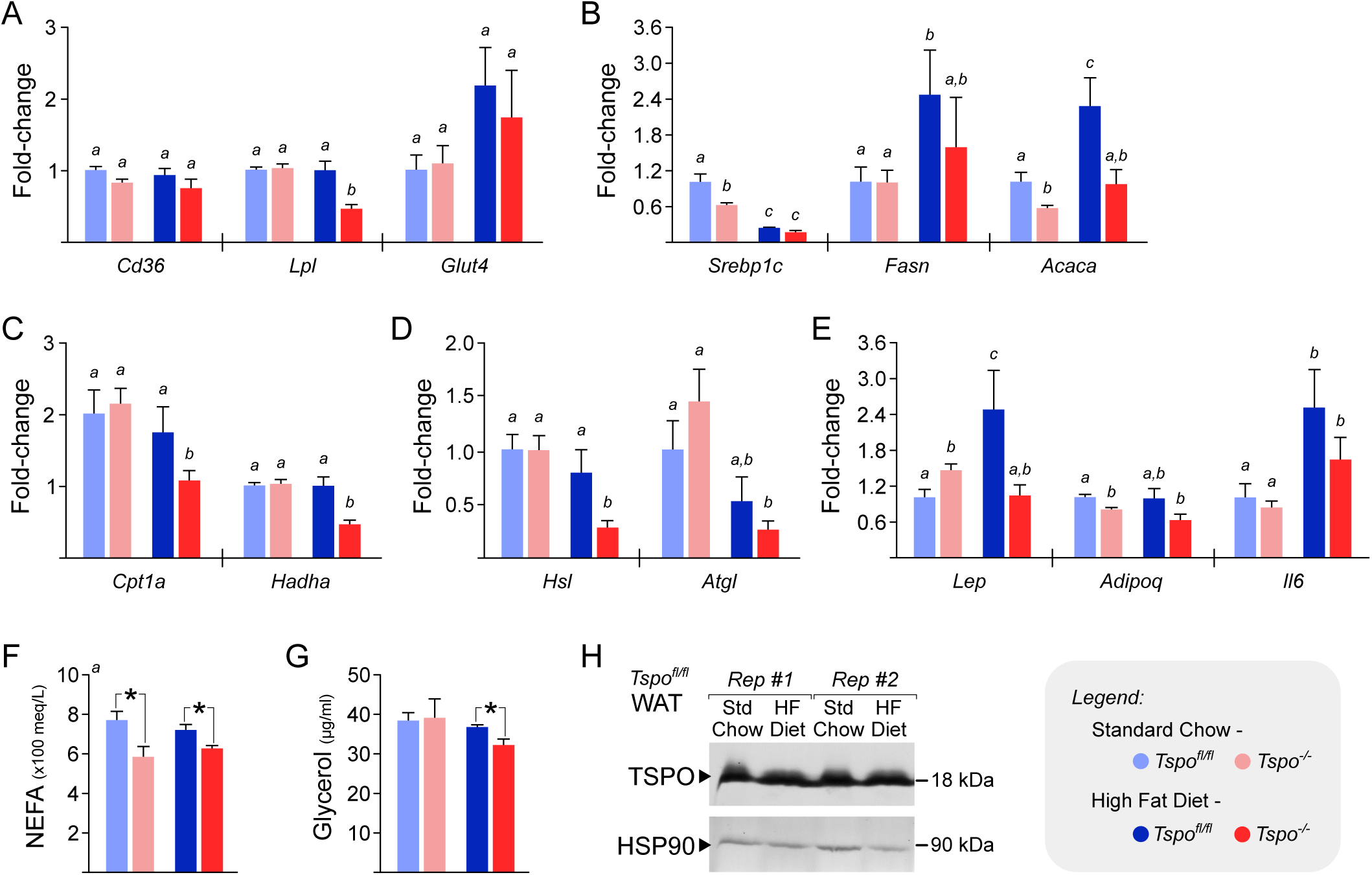
TSPO deficiency alters lipid metabolism gene expression in white adipose tissue following high-fat diet feeding. (A) Relative mRNA expression of lipid uptake and glucose transport genes (*Cd36*, *Lpl*, *Glut4*) in gonadal white adipose tissue (WAT). Expression of *Lpl* was significantly downregulated in *Tspo^-/-^* mice after 8 weeks of HF diet feeding, whereas *Cd36* and *Glut4* levels were not significantly altered based on genotype. (B) Relative mRNA expression of lipogenesis-related genes (*Srebp1c*, *Fasn*, *Acaca*). Expression of *Acaca* was significantly downregulated in *Tspo^-/-^* mice after 8 weeks of HF diet feeding, whereas *Srebp1c* and *Fasn* were not significantly altered based on genotype. (C) Relative mRNA expression of fatty acid oxidation genes (*Cpt1a*, *Hadha*) in WAT. Expression of *Cpt1a* and *Hadha* significantly downregulated in *Tspo^-/-^* mice after 8 weeks of HF diet feeding. (D) Relative mRNA expression of the lipolytic enzymes (*Hsl* and *Atgl*) in WAT. Expression of *Hsl* was significantly downregulated in *Tspo^-/-^* mice after 8 weeks of HF diet feeding, whereas *Atgl* showed a decreasing trend that was not statistically significant. (E) Relative mRNA expression of adipokine genes (*Lep*, *Adipoq* and *Il6*) revealed significant downregulation of *Lep* in *Tspo^-/-^* mice after 8 weeks of HF diet feeding, whereas *Adipoq* and *Il6* were not significantly altered based on genotype. (F) Circulating non-esterified fatty acid (NEFA) concentrations measured in serum showed significant downregulation in *Tspo^-/-^* mice on both standard chow and HF diets. (G) Circulating serum glycerol concentrations also showed significant downregulation in *Tspo^-/-^* mice after 8 weeks of HF diet feeding. (H) Western blot analysis of TSPO and HSP90 protein expression in WAT under standard chow and HF diet conditions did not show any notable differences. Data are presented as mean ± SEM. Different letters denote statistically significant differences among groups (p < 0.05).

However, after HF diet feeding, *Lpl* expression was significantly downregulated in *Tspo^-/-^* WAT, while *Cd36* and *Glut4* expression remained unchanged (Figure 3A). Among FA synthesis-related genes, chow-fed *Tspo^-/-^* mice showed significant downregulation of *Srebp1c* and *Acaca*, with no difference in *Fasn* expression compared to *Tspo^fl/fl^* cohorts (Figure 3B), indicating an altered lipogenic program. Following HF diet feeding, *Acaca* expression remained significantly lower in *Tspo^-/-^* WAT, but *Srebp1c* and *Fasn* levels were comparable to controls (Figure 3B). In the mitochondrial β-oxidation pathway (fatty acid oxidation, FAO), no significant differences were observed in the expression of *Cpt1a* or *Hadha* in chow-fed *Tspo^-/-^* mice (Figure 3C). In contrast, after HF diet feeding, *Tspo^-/-^* WAT exhibited significant downregulation of both *Cpt1a* and *Hadha* (Figure 3C), indicating diminished FAO in *Tspo^-/-^* WAT. For genes regulating lipolysis, expression of *Hsl* and *Atgl* was not significantly altered by TSPO deficiency under chow diet conditions (Figure 3D). After HF diet feeding, however, *Tspo^-/-^* WAT showed a significant decrease in *Hsl* expression and a downward trend in *Atgl* expression (Figure 3D), indicating reduced lipolysis in *Tspo^-/-^* WAT. Finally, among adipokine genes, chow-fed *Tspo^-/-^* mice exhibited reduced *Adipoq* expression and elevated *Lep* expression, with no difference in *Il6* levels (Figure 3E). Under HF diet conditions, expression of *Adipoq* and *Il6* was not significantly different between genotypes, however, expression of *Lep* was significantly lower in *Tspo^-/-^* mice (Figure 3E). These findings demonstrate that while TSPO deficiency causes selective, inconsistent alterations in lipid metabolism gene expression under chow diet, it leads to more pronounced and coordinated changes following HF diet feeding.

### TSPO deficiency reduces circulating NEFA and glycerol

To confirm a functional impact on lipolysis, we measured circulating levels of non-esterified fatty acids (NEFA) and glycerol, key products of triglyceride breakdown. Consistent with reduced lipolytic gene expression, *Tspo^-/-^* mice exhibited significantly lower plasma NEFA concentrations compared to *Tspo^fl/fl^* controls under both chow and HF diet conditions (Figure 3F). Plasma glycerol concentrations were also significantly reduced in *Tspo^-/-^* mice following HF diet feeding, although no significant difference was observed under chow diet (Figure 3G). Finally, to determine whether HF diet influenced TSPO expression itself, we assessed TSPO protein levels in WAT by Western blot analysis and found no appreciable differences between chow-fed and HF diet-fed *Tspo^fl/fl^* mice (Figure 3H). These data indicate that TSPO deficiency lowers circulating lipolytic products.

### Lipidomic profiling reveals no major impact of TSPO deficiency on gWAT lipid composition

To assess whether TSPO deficiency alters lipid composition in white adipose tissue, we performed targeted lipidomic analysis of gWAT. A total of 71 annotated lipid species were detected across four major classes: free fatty acids (FAs), diacylglycerols (DGs), triacylglycerols (TGs), and cholesterol. Heatmap visualization revealed diet-dependent clustering, with HF diet samples exhibited some distinctions among the different lipids relative to chow-fed controls, regardless of TSPO genotype (Figure 4). Among FAs, species such as FA(18:3) was consistently lower in the HF diet in both genotypes. Among DGs, species such as DG(34:1) and DG(36:2) were consistently elevated in the HF diet in both genotypes. Among TGs, species such as TG(56:7), and TG(58:10) were consistently lower in both *Tspo^fl/fl^* and *Tspo^-/-^* mice under HF diet. Regardless, no consistent genotype-specific differences were observed within either dietary group with minor variability across individual samples but no evidence of systematic shifts due to TSPO deletion. These data demonstrate that TSPO deficiency does not appreciably affect the abundance or distribution of major lipid species in gWAT, and that lipid composition changes with high-fat diet are preserved in the absence of TSPO.

**Figure 4.**
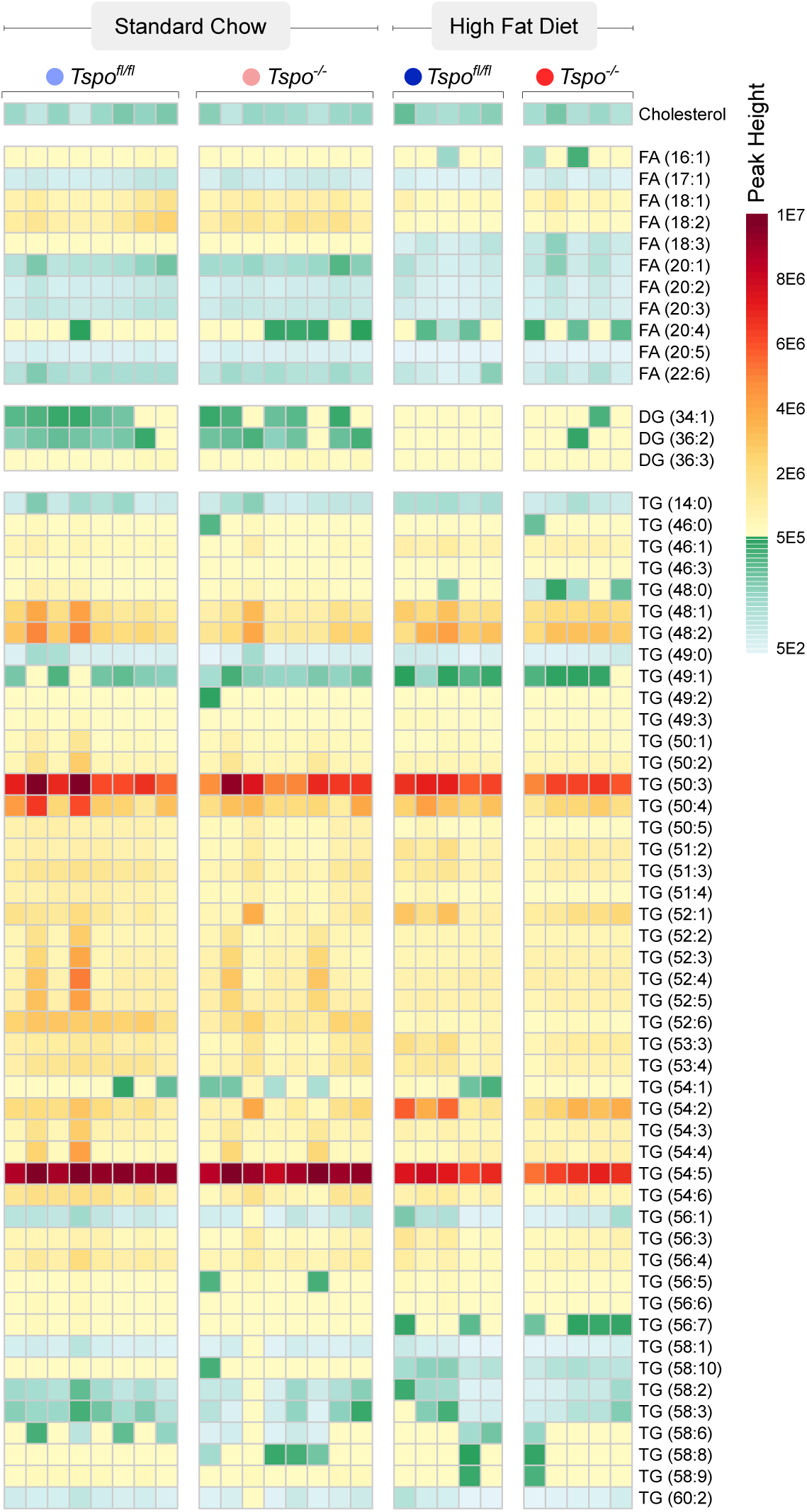
TSPO deficiency does not alter lipid composition in gWAT at baseline or after high-fat diet. Relative abundance of detected lipid species was quantified and visualized as a heatmap. Each row represents a lipid species categorized by lipid class: cholesterol, free fatty acids (FA), diacylglycerols (DG), and triacylglycerols (TG). Columns represent individual biological replicates from four groups: *Tspo^fl/fl^* mice and *Tspo^-/-^* mice on standard chow; *Tspo^fl/fl^* and *Tspo^-/-^* mice on 8 weeks of high-fat diet as indicated. Lipid signal intensity is color-coded on a dual scale (low range – from 500-5E5 peak height; high range from 5E5 to 1E7). Standard chow and high-fat diet conditions exhibit some minor distinctions patterns in lipid abundance, but there were no notable compositional differences between gWAT from *Tspo^fl/fl^* mice and *Tspo^-/-^* mice.

### TSPO deficiency enhances lipid accumulation in primary preadipocyte diDerentiation

To determine whether TSPO deficiency affects adipocyte differentiation and lipid metabolism *in vitro*, we compared lipid accumulation, protein expression, and metabolic gene profiles in primary preadipocytes isolated from *Tspo^fl/fl^* and *Tspo^-/-^* mice. Oil Red O staining at Day 4 of differentiation revealed greater lipid accumulation in *Tspo^-/-^* adipocytes compared to *Tspo^fl/fl^* controls (Figure 5A), a finding confirmed by quantification of Oil Red O absorbance (Figure 5B). Quantitative PCR analysis in *Tspo^fl/fl^* adipocytes showed progressive induction of *Tspo* mRNA expression over the course of differentiation, consistent with increased TSPO expression during adipogenesis (Figure 5C). Western blot analysis demonstrated an expected increase in PPARγ protein levels during differentiation, confirming successful adipogenic induction; TSPO protein levels were also elevated over the course of differentiation in *Tspo^fl/fl^* cells, while HSP90 levels remained constant across time points (Figure 5C). Increase in TSPO expression is consistent with previous reports correlating increasing TSPO levels and adipogenesis (Wade *et al*. 2005). Additional Western blotting of Day 4 adipocytes from independent replicate samples demonstrated that PPARγ and FABP4 protein levels were reduced in *Tspo^-/-^* adipocytes relative to *Tspo^fl/fl^* controls (Figure 5E), suggesting impaired adipogenic programming.

**Figure 5.**
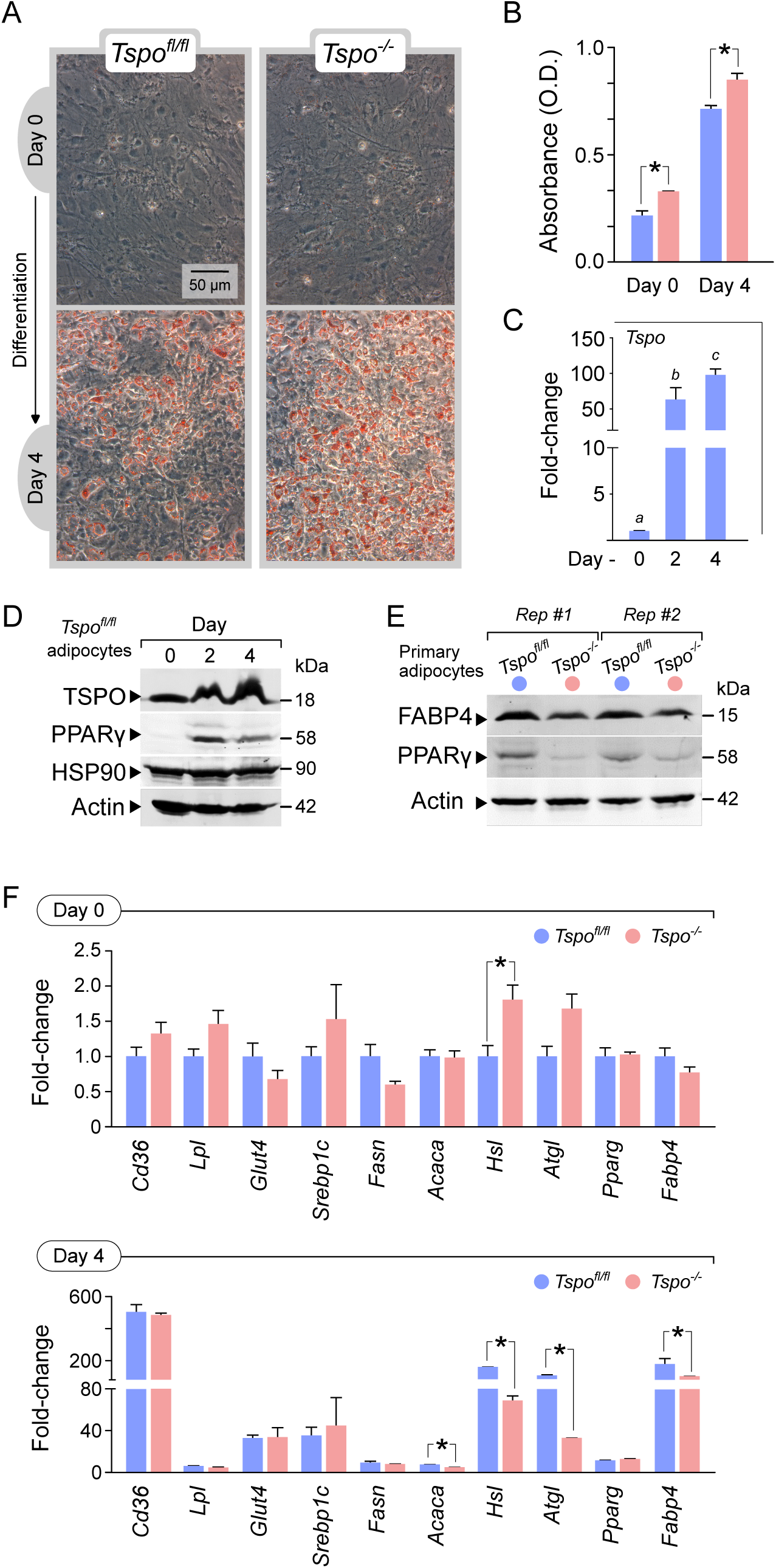
TSPO deficiency enhances lipid accumulation in primary preadipocytes diTerentiation. (A) Representative Oil Red O staining of primary adipocytes derived from *Tspo^fl/fl^* and *Tspo^-/-^* mice at Day 4 of differentiation under adipogenic conditions indicated higher abundance of lipid accumulation in *Tspo^-/-^* adipocytes. (B) Graph showing quantification of lipid accumulation by Oil Red O absorbance at Day 0 and Day 4 of differentiation. *Tspo^-/-^* adipocytes showed significantly higher levels of lipid accumulation compared to *Tspo^fl/fl^* controls. (C) Quantitative PCR analysis of *Tspo* mRNA expression during differentiation (Days 0, 2, and 4) in *Tspo^fl/fl^* adipocytes demonstrates robust induction of Tspo transcript levels over the course of adipogenesis. (D) Western blot analysis of TSPO, PPARγ, and HSP90 protein levels at Days 0, 2, and 4 of differentiation in *Tspo^fl/fl^* adipocytes indicated upregulation of TSPO and PPARγ at Day 2 and 4; HSP90 levels remained unchanged with differentiation. Actin is shown as a loading control. (E) Western blot analysis of PPARγ and FABP4 protein expression in Day 4 adipocytes from two independent replicate samples of *Tspo^fl/fl^* and *Tspo^-/-^* mice. *Tspo^-/-^* cells show reduced PPARγ and FABP4 protein levels relative to controls. Actin serves as a loading control. (F) Relative mRNA expression of adipogenesis-associated genes (*Pparg, Fabp4, Cd36, Lpl, Glut4, Srebp1c, Fasn, Acaca, Hsl, Atgl*) in *Tspo^fl/fl^* and *Tspo^-/-^* adipocytes at Days 0 and 4. At Day 4, *Tspo^-/-^* adipocytes exhibit significant downregulation of *Fabp4*, *Acaca*, *Hsl*, and *Atgl*, consistent with impaired adipogenic and lipolytic gene programs (mean ± SEM; *p < 0.05).

To further characterize the molecular basis of these phenotypic changes, we assessed the expression of key genes involved in nutrient uptake, fatty acid metabolism, and adipogenesis at Days 0 and 4 of differentiation. At Day 0, undifferentiated *Tspo^-/-^* preadipocytes exhibited significant upregulation of *Hsl* (Figure 5F). At Day 4, differentiated *Tspo^-/-^* adipocytes exhibited significant downregulation of *Fabp4*, *Acaca*, and both major lipolytic enzymes *Hsl* and *Atgl* compared to *Tspo^fl/fl^* controls (Figure 5F). Expression of other nutrient transport and adipogenic markers, remained largely unchanged between genotypes (Figure 5F). These findings indicate that TSPO deficiency impairs adipocyte lipid metabolism by suppressing key components of the lipolytic pathway during differentiation, consistent with the defects observed in adipose tissue *in vivo*.

### Pharmacological modulation of TSPO impacts adipocyte diDerentiation

To assess whether pharmacological modulation of TSPO affects adipocyte differentiation, we treated primary adipocytes derived from *Tspo^fl/fl^* and *Tspo^-/-^* mice with the TSPO-binding drugs PK11195 or etifoxine. As shown by Oil Red O staining at day 4, *Tspo^-/-^* adipocytes accumulated significantly more lipid than *Tspo^fl/fl^* controls under basal conditions, consistent with previous observations (Figure 6A). Notably, treatment of *Tspo^fl/fl^* adipocytes with either PK11195 or etifoxine increased lipid accumulation to levels comparable to untreated *Tspo^-/-^* cells (Figure 6A-B), suggesting that TSPO-binding drugs phenocopy the knockout state. Supporting this interpretation, PK11195 and etifoxine treatment had no enhancing effect on the already higher level of lipid accumulation observed in *Tspo^-/-^* adipocytes (Figure 6B).

**Figure 6.**
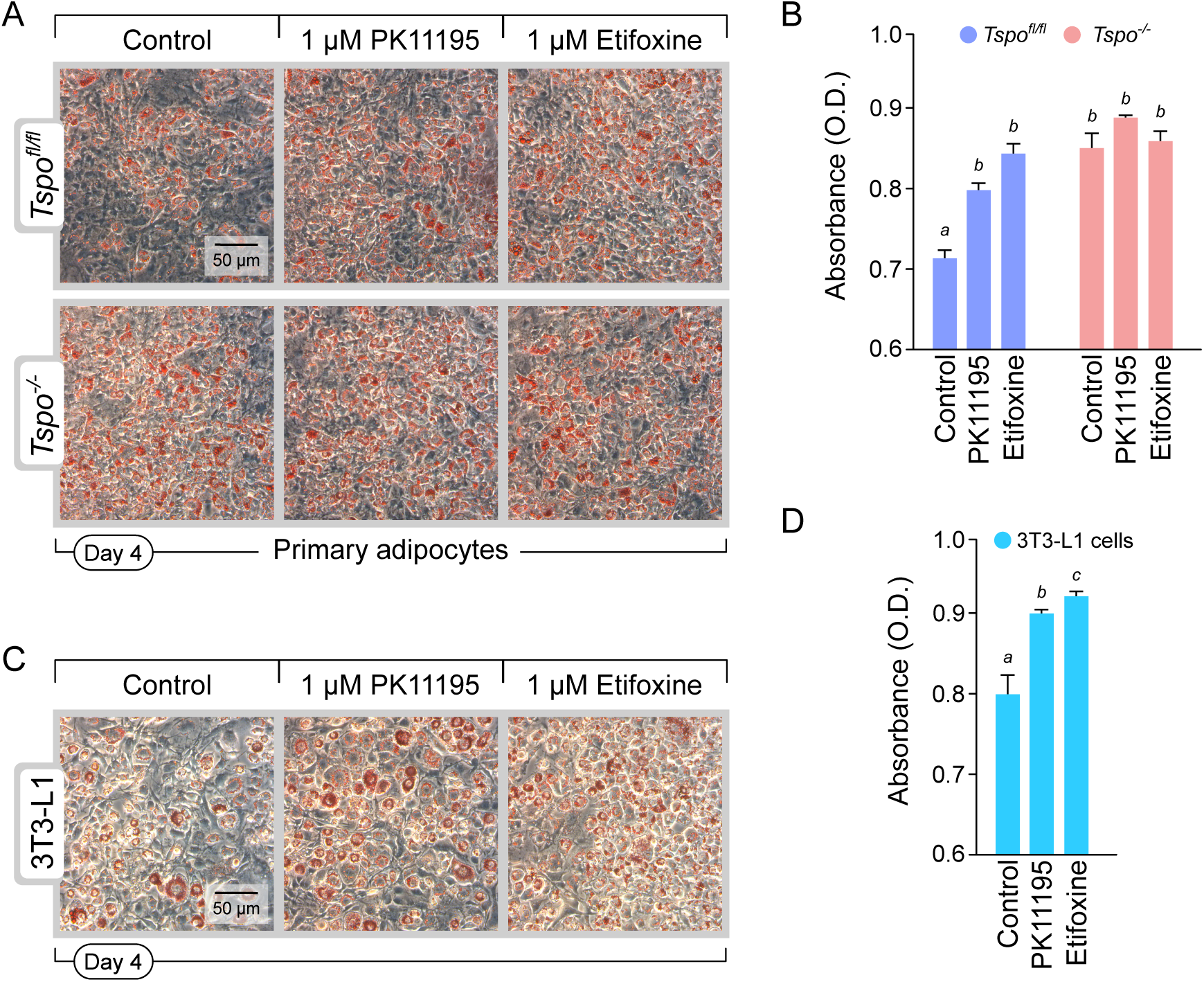
Pharmacological modulation of TSPO impairs adipocyte diTerentiation. (A) Representative Oil Red O staining of primary adipocytes derived from *Tspo^fl/fl^* and *Tspo^-/-^* mice treated during differentiation with vehicle control, PK11195 (1 μM), or etifoxine (1 μM). Higher levels of lipid accumulation could be observed in *Tspo^fl/fl^* adipocytes with PK11195 and etifoxine treatments. (B) Quantification of lipid accumulation by Oil Red O absorbance (520 nm) in primary adipocytes, showed increased lipid accumulation following PK11195 and etifoxine treatments in *Tspo^fl/fl^* adipocytes to levels that were comparable to *Tspo^-/-^* adipocytes. PK11195 and etifoxine had no effect on the already significantly higher levels of lipid accumulation seen in *Tspo^-/-^* adipocytes. (C) Representative Oil Red O staining of 3T3-L1 preadipocytes differentiated in the presence of vehicle control, PK11195, or Etifoxine. Higher levels of lipid accumulation could be observed in 3T3-L1 adipocytes with PK11195 and etifoxine treatments. (D) Quantification of lipid accumulation by Oil Red O absorbance in 3T3-L1 adipocytes, showed increased lipid accumulation following PK11195 and etifoxine treatments. Data are presented as mean ± SEM. Different letters denote statistically significant differences among groups (p < 0.05).

To test whether this pharmacological response was conserved in an established cell line, 3T3-L1 preadipocytes were treated with PK11195 or etifoxine under standard differentiation conditions. Both compounds significantly increased lipid accumulation (Figure 6C-D), consistent with findings in *Tspo^fl/fl^* primary cells. These findings demonstrate that pharmacological binding of TSPO enhances adipogenesis in a manner that mimics genetic deletion of the protein.

### TSPO regulates porphyrin homeostasis

To investigate whether TSPO influences porphyrin and heme biosynthesis during adipogenesis, we first quantified protoporphyrin IX (PPIX) concentrations and the expression of key porphyrin and heme biosynthetic enzymes in WAT. At baseline, PPIX levels in WAT were comparable between *Tspo^fl/fl^* and *Tspo^-/-^* mice (Figure 7A). However, under HF diet conditions, Tspo*^-/-^* mice exhibited significantly reduced expression of *Alas* (δ-aminolevulinic acid synthase) and *Fech* (ferrochelatase), the first and last enzymes in the heme biosynthesis pathway, respectively, relative to *Tspo^fl/fl^* controls (Figure 7B).

**Figure 7.**
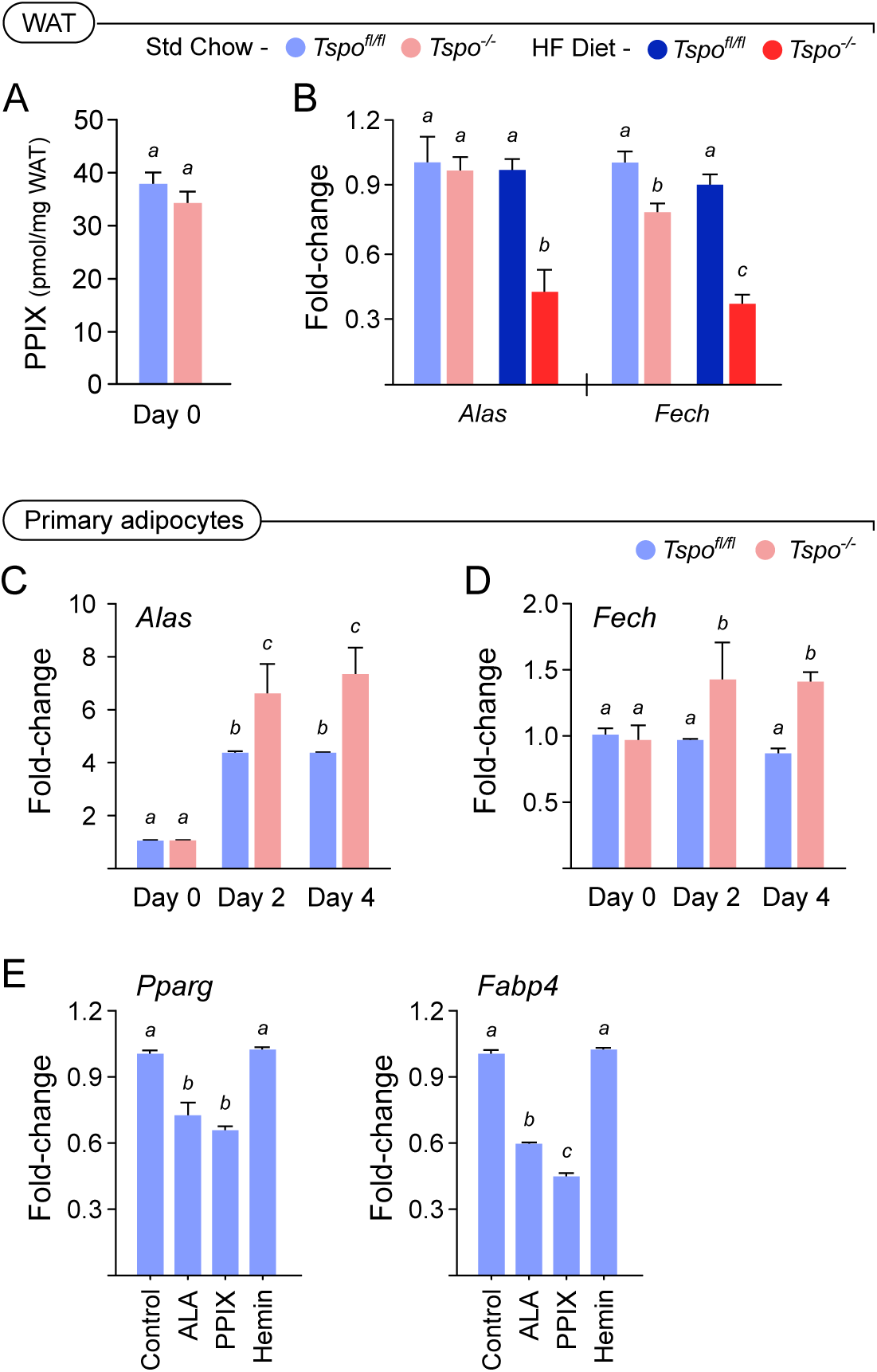
TSPO deficiency alters porphyrin and heme biosynthesis gene expression in white adipose tissue and during adipocyte diTerentiation. (A) Baseline protoporphyrin IX (PPIX) concentrations measured in WAT was not different between *Tspo^fl/fl^* and *Tspo^-/-^* mice under standard chow conditions. (B) Relative mRNA expression of δ-aminolevulinic acid synthase (*Alas*) and ferrochelatase (*Fech*) in WAT under HF diet conditions showed significant downregulation in *Tspo^-/-^* mice compared to *Tspo^fl/fl^* controls. (C-D) During *in vitro* adipocyte differentiation, *Tspo^-/-^* primary adipocytes showed significantly elevated expression of *Alas* and *Fech* at days 2 and 4 compared to *Tspo^fl/fl^* cells, suggesting dysregulation of porphyrin and heme biosynthetic genes in the absence of TSPO. (E) Adipogenic gene expression in *Tspo^fl/fl^* primary adipocytes treated with δ-aminolevulinic acid (ALA), protoporphyrin IX (PPIX), or hemin during differentiation at Day 4. Treatment with ALA and PPIX significantly reduced mRNA levels of the adipogenic markers *Pparg* and *Fabp4*, whereas hemin had no significant effect. Data are presented as mean ± SEM. Different letters denote statistically significant differences among groups (p < 0.05).

During *in vitro* adipocyte differentiation of primary adipocytes from *Tspo^fl/fl^* and *Tspo^-/-^* mice, expression of both *Alas* and *Fech* was markedly upregulated in *Tspo^-/-^* primary adipocytes compared to *Tspo^fl/fl^* cells, with significant differences observed at both days 2 and 4 of differentiation (Figure 7C-D). These data suggest that TSPO deficiency leads to alterations in the regulation of porphyrin and possibly heme biosynthesis.

To assess whether modulation of porphyrin pathway intermediates directly impacts adipogenic gene expression, we treated *Tspo^fl/fl^* primary preadipocytes with δ-aminolevulinic acid (ALA), PPIX, or hemin during differentiation. Both ALA and PPIX at Day 4 significantly suppressed expression of the adipogenic markers *Pparg* and *Fabp4*, whereas hemin had no effect on their expression (Figure 7E). These findings indicate that the accumulation of upstream porphyrin intermediates negatively regulates adipocyte differentiation, and that TSPO may influence adipogenesis in part by modulating porphyrin levels.

### PPIX accumulation inhibits adipocyte diDerentiation and lipolytic responses

To investigate whether porphyrin biosynthetic intermediates modulate adipocyte function, we first measured intracellular PPIX levels during 3T3-L1 adipocyte differentiation. PPIX levels increased progressively from day 0 to day 4, indicating enhanced flux through the porphyrin synthesis pathway during adipogenesis (Figure 8A). Consistent with this, *Alas* mRNA, encoding the rate-limiting enzyme in porphyrin biosynthesis, was significantly upregulated during differentiation. In contrast, *Fech*, which catalyzes the terminal step converting PPIX to heme, showed a declining expression pattern (Figure 8B), suggesting a potential accumulation of porphyrin intermediates.

**Figure 8.**
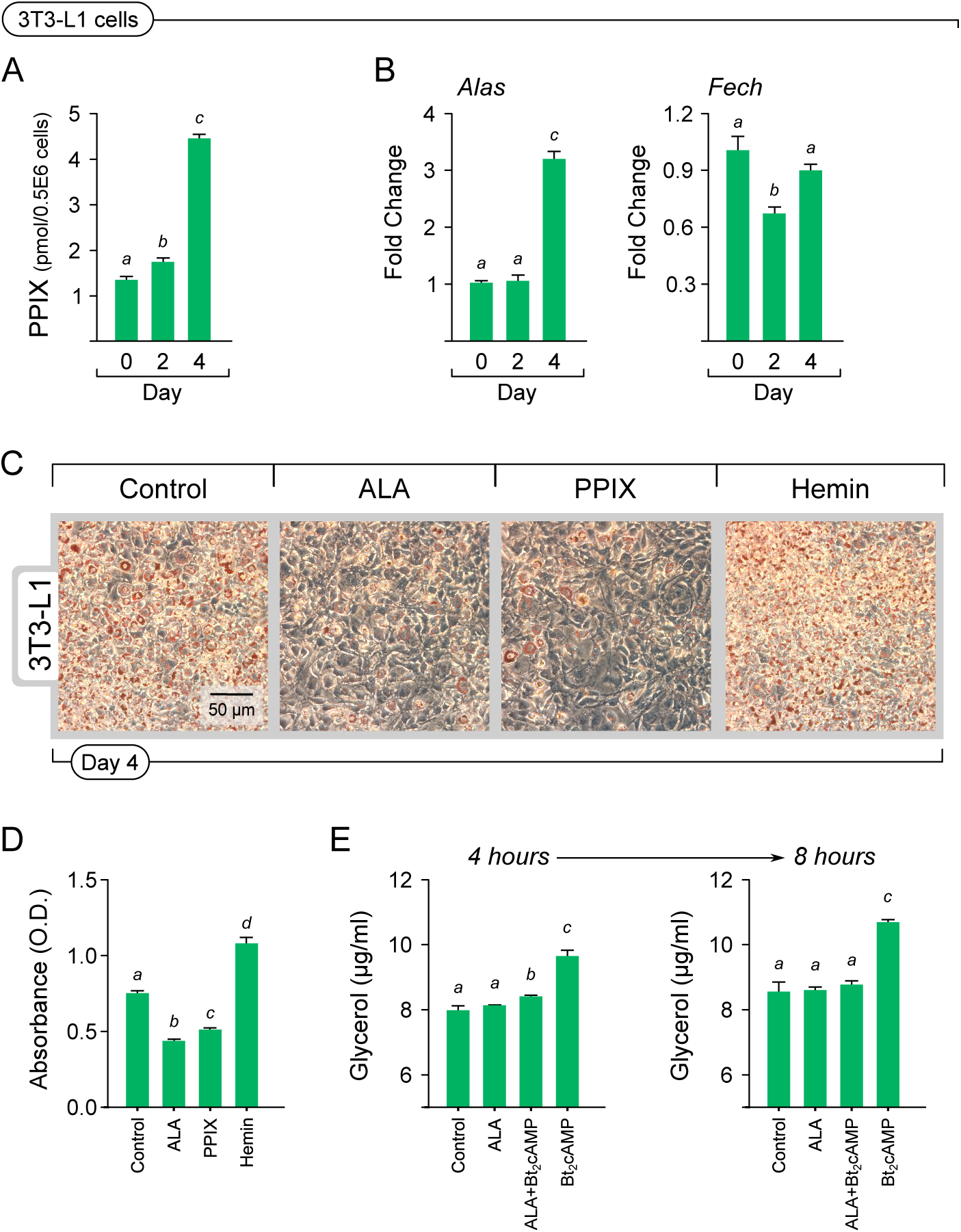
Porphyrin intermediates inhibit adipocyte diTerentiation and lipolytic responses. (A) Cellular protoporphyrin IX (PPIX) levels increase progressively during 3T3-L1 differentiation, linking porphyrin biosynthesis to adipogenic progression. (B) *Alas* and *Fech* mRNA expression during differentiation revealed a significant upregulation of *Alas* by Day 4, and downregulation of *Fech* by Day 2, indicating activation of porphyrin synthesis with reduced terminal heme synthesis. (C-D) Representative Oil Red O staining and quantification of lipid accumulation in 3T3-L1 cells treated with ALA (1 mM), PPIX (1 µM), or hemin (10 µM) at Day 4 of differentiation. ALA and PPIX significantly reduced lipid accumulation, whereas hemin promoted it, suggesting that upstream porphyrins, but not heme, suppress adipogenesis. (E) Glycerol release at 4-and 8-hours post-treatment with ALA (1 mM), ALA + Bt_2_cAMP (0.5 mM), or Bt_2_cAMP alone showed that ALA significantly suppressed cAMP-stimulated lipolysis. Bars represent mean ± SEM; different letters indicate statistically significant differences (p < 0.05).

To test the functional significance of porphyrin intermediates, 3T3-L1 cells were treated with δ-aminolevulinic acid (ALA), PPIX, or hemin from day 0 to day 4 of differentiation. Oil Red O staining revealed that both ALA and PPIX treatments significantly reduced lipid accumulation compared to control, whereas hemin treatment had the opposite response significantly increasing lipid accumulation in differentiated 3T3-L1 cells (Figure 8C). These results indicated that PPIX, but not heme, impairs adipocyte differentiation.

We next assessed the impact of porphyrin intermediates on lipolytic responses in mature adipocytes. Glycerol release assays demonstrated that ALA treatment suppressed both basal and cAMP-stimulated lipolysis at 4-and 8-hours post-treatment (Figure 8D), indicating that porphyrin accumulation also negatively regulates lipolytic pathways. These findings suggest that dysregulation or accumulation of porphyrin intermediates and PPIX selectively impairs both adipogenic and lipolytic programs in adipocytes, independent of terminal heme synthesis.

## Discussion

The long-held assumption that the translocator protein (TSPO) facilitates mitochondrial cholesterol import for steroidogenesis has been unequivocally overturned by genetic studies showing normal hormone biosynthesis in TSPO knockout models (Morohaku *et al*. 2014; Tu *et al*. 2014, 2015). However, an explanation for high TSPO expression in both steroidogenic and non-steroidogenic cells that are metabolically active in lipid turnover remains unclear. Our previous work demonstrated that TSPO deletion in Leydig cells leads to a metabolic reprogramming favoring mitochondrial fatty acid oxidation (FAO), despite unaltered steroidogenic output (Tu *et al*. 2016). In parallel, we reported that TSPO expression in hamster adrenals is tightly linked to triglyceride handling rather than hormone biosynthesis, implicating a broader role for TSPO in lipid metabolic regulation (Koganti & Selvaraj 2020). The current study expands this framework to adipose tissue, revealing that TSPO loss in white adipocytes disrupts lipolytic capacity and mitochondrial FAO, leading to neutral lipid accumulation, impaired fatty acid mobilization, and a distinctive porphyrin-sensitive metabolic phenotype. Together, these findings reposition TSPO as a mitochondrial integrator of lipid flux through porphyrins, with physiological relevance across diverse metabolic tissues.

We found no apparent metabolic perturbations at baseline in *Tspo^-/-^* mice, consistent with previous reports showing normal body weight, food intake, and adiposity under standard chow-fed conditions (Morrissey *et al*. 2021). However, exposure to HF diet unmasked a robust adipose phenotype characterized by increased WAT mass and adipocyte hypertrophy, despite unchanged feed intake in *Tspo^-/-^* mice. This expansion was not attributable to enhanced nutrient uptake or lipogenesis, as expression of *Cd36*, *Glut4*, *Fasn*, and *Srebp1c* remained unchanged or was suppressed. Instead, the phenotype reflected a failure in lipid clearance: expression of the canonical lipolytic enzymes *Hsl* and *Atgl* was significantly reduced, accompanied by a decrease in *Lpl* and circulating glycerol and non-esterified fatty acids. Notably, NEFA levels were already significantly lower in chow-fed *Tspo^-/-^* mice, suggesting a suppression of basal lipolysis that becomes exacerbated under conditions of metabolic stress. These data point to a cell-autonomous deficit in triglyceride mobilization. Particularly, this impairment extended to the transcriptional suppression of mitochondrial FAO enzymes (*Cpt1a*, *Hadha*), suggesting that the absence of TSPO compromises the coordinated coupling of lipolysis and downstream fatty acid oxidation.

Importantly, lipidomic profiling of gWAT revealed that neither baseline lipid composition nor HFD-induced lipid species enrichment was significantly altered by TSPO deficiency. High-fat feeding robustly increased long-chain and unsaturated triacylglycerols across genotypes, indicating that the observed adipocyte lipid accumulation in *Tspo^-/-^* mice is not due to aberrant lipid uptake or storage but rather a failure to mobilize lipids already present. This decoupling is especially evident under conditions of metabolic overload, where intact TSPO appears necessary to maintain lipid flux and prevent excessive lipid accumulation in adipocytes.

A striking mechanistic insight from this study is the identification of PPIX, a long known endogenous TSPO ligand (Verma *et al*. 1987), and intermediate in the heme biosynthesis pathway, as a negative regulator of adipocyte lipid metabolism. During adipogenesis, intracellular PPIX levels rise, and we found that providing PPIX or its precursor ALA in excess, selectively suppressed adipogenic and lipolytic gene expression, while hemin, representing the terminal product of the pathway, had no such effect. These results implicate porphyrin intermediates, rather than heme, as bioactive modulators of adipocyte function. Notably, *Tspo* transcript and protein levels increased during adipocyte differentiation, consistent with prior reports demonstrating TSPO upregulation in response to adipogenic cues (Wade *et al*. 2005), suggesting that enhanced TSPO expression may serve a functional role in buffering or regulating porphyrin accumulation during this transition.

The loss of TSPO altered the expression dynamics of porphyrin biosynthetic enzymes: *Alas*, the first and rate-limiting enzyme of the porphyrin biosynthetic pathway (Hunter & Ferreira 2009), and *Fech*, which catalyzes the conversion of PPIX to heme (Lecerof *et al*. 2000), were diminished *in vivo* under HFD, whereas both *Alas* and *Fech* were transcriptionally upregulated during *in vitro* differentiation. This suggests that TSPO acts to constrain the accumulation or signaling activity of porphyrin intermediates, possibly through binding, sequestration, or facilitating their mitochondrial processing (Wendler *et al*. 2003; Vanhee *et al*. 2011; Busch *et al*. 2017). In its absence, dysregulated porphyrin metabolism may impair mitochondrial function and directly suppress the transcriptional programs required for lipid mobilization. This TSPO–PPIX axis provides a previously unrecognized mechanism by which mitochondrial tetrapyrrole metabolism intersects with adipocyte bioenergetics and lipid turnover.

Our use of the TSPO-binding drugs PK11195 and etifoxine provided a critical pharmacological lens through which to validate the metabolic functions of TSPO identified in the genetic model. Both ligands recapitulated the key phenotype of TSPO deficiency namely, enhanced lipid accumulation in *Tspo^fl/fl^* primary adipocyte. Importantly, there was no effect on *Tspo^-/-^* primary adipocytes, confirming that these actions are mediated specifically through TSPO rather than ob-target effects. Nonetheless, it is important to recognize that both PK11195 (Tu *et al*. 2015) and etifoxine (do Rego *et al*. 2015) are known to exhibit additional pharmacological activities. Both PK11195, an isoquinoline carboxamide, widely considered the prototypical non-benzodiazepine TSPO-binding drug (Anholt *et al*. 1985), and etifoxine, a clinically used anxiolytic that binds TSPO with a distinct pharmacology (Boissier *et al*. 1972; Costa *et al*. 2017), similarly disrupted adipocyte lipid metabolism in our study, supporting a conserved mechanism of TSPO-dependent regulation (Fonia *et al*. 1996). Given the complexity of TSPO-drug interactions, which may be modulated by the lipid environment, and ligand-specific conformational shifts (Jaremko *et al*. 2014), our findings argue for a cautious interpretation of TSPO-targeted interventions in both experimental and clinical settings. The ability of these ligands to mimic genetic deletion effects also opens the possibility that specific ligand classes may differentially bias the TSPO-PPIX axis toward lipid-retentive or lipid-mobilizing states, offering a potential framework for the development of next-generation TSPO modulators with tailored metabolic effects.

The emerging paradigm from this and related studies is that TSPO modulates mitochondrial lipid metabolism through porphyrins as regulatory intermediates that are evolutionarily conserved. Cross-species investigations support this broader view: in plants, TSPO modulates tetrapyrrole regulation under stress conditions (Vanhee *et al*. 2011), and in cyanobacteria, TSPO binds diverse porphyrins and may act as a control for tetrapyrrole metabolism (Busch *et al*. 2017). In mammalian systems, TSPO loss has been linked to altered mitochondrial respiration and shifts in substrate utilization in microglia (Milenkovic *et al*. 2019), macrophages (Taylor *et al*. 2014), and retinal pigment epithelial cells (Alamri *et al*. 2019), many of which demonstrate compensatory reliance on glycolysis or disrupted FAO. Our findings in WAT extend these observations by showing that TSPO deficiency impairs the lipolysis and FAO not through overt mitochondrial dysfunction, but through disruption of PPIX-based metabolic coordination. Recent studies have indicated that TSPO’s affinity for PPIX might be coupled to conformational transitions (Yeh et al., 2023) and catalysis (Ginter *et al*. 2013), providing a plausible molecular basis for porphyrin-dependent regulation of mitochondrial processes. Although recent studies have taken steps to define TSPO as a PPIX oxygenase (Qiu *et al*. 2025), the activity is observed only under photoactivated conditions, with no evidence of an enzymatic function in the absence of light or native mitochondrial conditions to be relevant for mammalian *in vivo* systems. These insights, combined with functional studies showing that TSPO can ameliorate PPIX pathogenesis (Zeno *et al*. 2012; Rosenberg *et al*. 2013), support the model that TSPO serves as a critical interface between lipid mobilization, oxidative metabolism, and regulation of tetrapyrrole levels.

Taken together, our findings indicate that TSPO is a homeostatic modulator of PPIX regulation of lipid metabolism in adipocytes, acting at the intersection of mitochondrial fatty acid oxidation, and lipolysis. Building on our prior work in Leydig cells and adrenal tissues, revealing that PPIX, an endogenous TSPO ligand, suppresses both adipogenic and lipolytic programs, we uncover a novel feedback mechanism in which mitochondrial tetrapyrrole flux influences lipid turnover. The capacity of TSPO to modulate this axis, and the ability of its pharmacological ligands to phenocopy its deficiency, redefine the functional landscape of this enigmatic protein. These insights carry substantial implications for interpreting TSPO biology across disciplines, from neuroinflammation to metabolic disease, and raise the prospect that tissue-specific TSPO activity may govern discrete metabolic control through differential porphyrin flux. By reframing TSPO as a lipid-metabolic integrator, this work opens new directions for targeting mitochondrial dysfunction in obesity, and metabolic syndrome.

## Materials and methods

### Mice

The generation of *Tspo^fl/fl^* and *Tspo^-/-^* mice has been previously described (Morohaku *et al*. 2014; Tu *et al*. 2014). In treatment groups, 6-week-old mice were provided with either Teklad Global standard chow diet (18% Protein Rodent Diet; Envigo) or Teklad Diets commercial 45% high-fat diet (TD.06415; Envigo) for a period of 8 weeks. Feed intake and body weight were measured every 2 days. Both male and female mice were fed HF diet from 6 weeks to 14 weeks of age. Inguinal WAT, gonadal WAT and liver were collected at 14 weeks of age for different measurements. *Tspo^fl/fl^* and *Tspo^-/-^* mice on standard chow diet were used for preadipocyte cultures. All animals were maintained in accordance with the National Institutes of Health Guide for the Care and Use of Laboratory Animals, and all procedures were approved by the Institutional Animal Care and Use Committee of Cornell University.

### Adipocyte morphometry

Gonadal white adipose tissue (gWAT) depots were collected from male mice at 14 weeks of age following high-fat diet feeding. Tissues were fixed in 10% neutral buffered formalin for 24 hours at room temperature, dehydrated through graded ethanol series, cleared in xylene, and embedded in paraffin. Serial sections were cut at a thickness of 4 µm using a rotary microtome and mounted on glass slides. Sections were stained with Mayer’s hematoxylin solution and eosin Y alcoholic solution (both from Sigma-Aldrich) following standard protocols (Morohaku *et al*. 2013). Stained sections were imaged using a Leica DM 1000 LED microscope equipped with an ICC50 HD digital camera. For quantification, contiguous fields were captured at 10x magnification, and adipocyte area and density were measured using ImageJ software (version 1.48v; National Institutes of Health). Individual adipocytes were manually outlined, and mean adipocyte area (μm²) and cell size distribution frequencies were calculated. A minimum of 100-150 cells per sample/mouse were analyzed to ensure statistical robustness.

### Preadipocyte isolation and culture

Primary preadipocytes were isolated from subcutaneous inguinal white adipose tissue (iWAT) as previously described (Hausman *et al*. 2008). Briefly, iWAT depots were harvested from groups of mice at 6-8 weeks of age, minced into small fragments, and digested in a solution containing collagenase from Clostridium histolyticum (Sigma-Aldrich) and 4-(2-hydroxyethyl)-1-piperazineethanesulfonic acid (HEPES; Sigma-Aldrich) in phosphate-buffered saline (PBS) at 37°C with constant shaking for 30 minutes. The resulting cell suspension was filtered through a 100-μm mesh to remove undigested tissue, and the stromal vascular fraction (SVF) containing preadipocytes was collected by centrifugation at 500 × g for 5 minutes. Pelleted cells were resuspended and plated in Dulbecco’s Modified Eagle’s Medium (DMEM; Sigma-Aldrich) supplemented with 10% fetal bovine serum (FBS), 1% penicillin-streptomycin (P/S), and 1% non-essential amino acids (NEAA) (hereafter referred to as DMEM+FBS). Cell proliferation was monitored for expansion and replated for differentiation experiments. Fibroblast-like murine 3T3-L1 preadipocytes (American Type Culture Collection, ATCC) were cultured under identical conditions in DMEM+FBS. All cells were maintained at 37°C in a humidified incubator with 5% CO₂.

### Adipocyte diDerentiation and pharmacological treatments

Preadipocytes (primary or 3T3-L1) were cultured at 37°C in a humidified incubator with 5% CO₂ in DMEM+FBS until they reached two days post-confluence (designated as Day 0). To induce adipogenic differentiation, preadipocytes were treated with differentiation medium consisting of DMEM+FBS supplemented with 0.5 mM 3-isobutyl-1-methylxanthine (IBMX; Sigma-Aldrich), 2.5 μM dexamethasone (Sigma-Aldrich), 10 μg/mL bovine insulin (Sigma-Aldrich), and 1 µM rosiglitazone (Sigma-Aldrich) for 2 days. After two days (Day 2), cells were switched to maintenance medium containing DMEM+FBS supplemented with 10 μg/mL bovine insulin alone. Maintenance medium was refreshed every day, and cells were cultured for an additional 2 days until fully differentiated, as indicated by extensive lipid droplet accumulation. For ligand treatment experiments, PK11195 (1 μM; Sigma-Aldrich), or etifoxine (1 μM; Sigma-Aldrich) was added to the culture medium beginning on Day 0 and replenished with each medium change until the completion of differentiation. At timepoints, cells were collected and processed for different measurements.

### Quantification of lipid accumulation

Primary preadipocytes and 3T3-L1 cells were cultured under adipogenic conditions and analyzed at Day 0 (undifferentiated) and Day 4 (fully differentiated) by staining for lipid accumulation as previously described (Koganti *et al*. 2022). Cells were gently washed twice with phosphate-buffered saline (PBS) and fixed with 4% paraformaldehyde (Electron Microscopy Sciences) for 15 minutes at room temperature. Following fixation, cells were washed sequentially with double-distilled water and 60% isopropanol to prepare for staining. Lipid droplets were visualized by staining with 0.3% Oil Red O (ORO) solution (Matheson, Coleman & Bell) for 30 minutes at room temperature. Excess stain was removed by washing with double-distilled water. For quantitative analysis, bound Oil Red O was eluted with 100% isopropanol (Mallinckrodt Chemicals), and absorbance was measured at 500 nm using an Infinite F50 plate reader (Tecan). Representative images of stained cells were acquired using a Leica DMIL LED inverted microscope equipped with an MC120 HD camera.

### Gene expression analysis

Total RNA was extracted from gWAT depots of male mice, primary adipocytes, and 3T3-L1 cells using TRIzol reagent (Life Technologies) according to the manufacturer’s instructions. RNA concentration and purity were assessed spectrophotometrically using a NanoDrop instrument (Thermo Fisher Scientific). Complementary DNA (cDNA) was synthesized from 1000 ng of total RNA using the High-capacity cDNA reverse transcription kit (Applied Biosystems), following the manufacturer’s protocol. Quantitative real-time PCR (qPCR) was performed using SYBR Green detection chemistry (Affymetrix) on a StepOne Plus Real-Time PCR System (Applied Biosystems). Primer sequences were obtained from the PrimerBank database (Harvard Medical School; listed in Table S1) and validated for specificity and amplification efficiency. Gene expression data were normalized to TATA-binding protein (Tbp) as the reference gene. For tissue samples, normalization was performed relative to *Tspo^fl/fl^* controls; for primary adipocytes, data were normalized to Day 0 *Tspo^fl/fl^* samples; and for 3T3-L1 cells, data were normalized to Day 0 vehicle-treated control samples. Relative expression levels were calculated using the comparative Ct (2^-ΔΔCt^) method (Livak & Schmittgen 2001).

### Western blot analysis

Western blotting was performed as previously described (Morohaku *et al*. 2013). Briefly, samples were lysed and processed in Laemmli sample buffer containing SDS and β-mercaptoethanol (Laemmli 1970). Protein concentrations were determined using a bicinchoninic acid (BCA) protein assay kit (Thermo Fisher Scientific) according to the manufacturer’s instructions. Equal amounts of protein (25 μg per sample) were separated by SDS-polyacrylamide gel electrophoresis (SDS-PAGE) and transferred onto polyvinylidene fluoride (PVDF) membranes (Millipore). Membranes were blocked for 1 hour at room temperature in 5% non-fat dry milk prepared in Tris-buffered saline containing 0.2% Tween-20 (TBS-T). Primary antibody incubations were performed overnight at 4°C using the following antibodies: rabbit anti-TSPO (EPR5384) monoclonal antibody (Abcam, #ab109497), rabbit anti-HSP90 (C45G5) monoclonal antibody (Cell Signaling, #4877), rabbit anti-PPARγ monoclonal antibody (Cell Signaling, #81B8), rabbit anti-FABP4 polyclonal antibody (Cell Signaling, #2120) and mouse monoclonal anti-β-actin antibody (Li-Cor, 926-42212) as a loading control. After washing with TBS-T, membranes were incubated with fluorescently labeled secondary antibodies: IRDye 800CW goat anti-rabbit IgG and IRDye 680RD goat anti-mouse IgG (Li-Cor). Fluorescent signals were detected and quantified using an Odyssey infrared imaging system (Li-Cor).

### Plasma NEFA quantification

Plasma non-esterified fatty acid (NEFA) concentrations were measured using the NEFA-HR(2) colorimetric assay kit (Wako Diagnostics) according to the manufacturer’s instructions. Blood was collected into EDTA-coated tubes, centrifuged at 1,500 × g for 10 minutes at 4°C, and the plasma supernatant was stored at -80°C until analysis. Plasma samples were thawed on ice, diluted appropriately with deionized water, and loaded into a 96-well microplate along with NEFA standards. Assay reagent A, containing coenzyme A, acyl-CoA synthetase, and acyl-CoA oxidase, was added to each well and incubated at 37°C for 5 minutes. Following this, assay reagent B, containing peroxidase and chromogenic substrates, was added, and plates were incubated for an additional 5 minutes at 37°C. Absorbance was measured at 550 nm using a microplate reader (Infinite F50, Tecan). NEFA concentrations were calculated based on a standard curve generated from known NEFA concentrations. All samples and standards were assayed in duplicate.

### Plasma glycerol quantification

Plasma glycerol concentrations were measured using a colorimetric enzymatic assay with the Free Glycerol Reagent (Sigma-Aldrich) according to the manufacturer’s instructions. Blood was collected into EDTA-coated tubes, centrifuged at 1,500 × g for 10 minutes at 4°C, and the plasma supernatant was stored at -80°C until analysis. Plasma samples were thawed on ice, diluted appropriately with deionized water, and loaded into a 96-well microplate along with glycerol standards. The Free Glycerol Reagent, containing glycerol kinase, glycerol phosphate oxidase, and peroxidase enzymes, was added to each well. Following incubation at 37°C for 5 minutes, absorbance was measured at 540 nm using a microplate reader (Infinite F50, Tecan). Glycerol concentrations were determined by comparison to a standard curve generated from known concentrations of glycerol. All samples and standards were assayed in duplicate.

### Lipidomics

Targeted lipidomics analysis of gWAT was performed using reverse-phase chromatography coupled with quadrupole time-of-flight mass spectrometry (CSH-QTOF MS), as previously described (Tu *et al*. 2017; Koganti *et al*. 2022), with minor modifications. Briefly, frozen gWAT samples were homogenized and extracted using a biphasic mixture composed of 225 µL cold methanol containing internal standards and 188 µL LC-MS grade water. Samples were vortexed and centrifuged at 14,000 x g for 2 minutes to separate phases. The upper organic layer (350 µL), enriched in lipids, was collected and dried using a vacuum centrifuge (Labconco). Dried lipid extracts were reconstituted in 110 µL of 90:10 methanol:toluene containing 50 ng/mL CUDA (Cayman Chemical) and analyzed using an Agilent 1290 Infinity LC system. Samples were analyzed in both positive and negative electrospray ionization (ESI) modes to broaden lipid coverage. For ESI(+) mode, mobile phases were modified with 10 mM ammonium formate and 0.1% formic acid, which enhanced detection of CE, DG, and PC classes. For ESI(−) mode, 10 mM ammonium acetate was used. Lipid separation was performed using an Acquity UPLC CSH C18 column (100 x 2.1 mm, 1.7 µm) with a matching VanGuard pre-column (5 x 2.1 mm, 1.7 µm) maintained at 65°C and a flow rate of 0.6 mL/min. The mobile phase consisted of 60:40 acetonitrile:water (A) and 90:10 isopropanol:acetonitrile (B), with a gradient elution profile optimized for broad lipid class resolution. Lipids were detected using an Agilent 6550 iFunnel QTOF mass spectrometer with jet stream ESI source. Acquisition was performed over an m/z range of 50–1700 at a rate of 2 spectra/s. QC included method blanks and pooled plasma standards. Data were processed using MS-DIAL software. Lipids were annotated using an in-house retention time and accurate mass library based on LipidBlast. The neutral lipids were further subcategorized into TG, DG, and total cholesterol. The percentage expression (peak height) of these subcategories was calculated relative to the total neutral lipid expression. All analyses and heatmap representations were performed using R (R Core Team. 2024).

### PPIX measurement

Concentrations of PPIX in WAT and cultured primary adipocytes were quantified using a modified fluorometric assay as previously described (Zhao *et al*. 2016). For tissue analysis, freshly harvested WAT was homogenized in 1% Triton X-100 in Tris-buffered saline (TBS) and centrifuged at 5,000 x g for 10 min at 4 °C. PPIX fluorescence was measured in the supernatant using a fluorescence spectrophotometer (Infinite 200, Tecan) with excitation at 400 nm and emission at 660 nm. For cultured cells, primary preadipocytes were harvested at the indicated time points (days 0, 2, and 4), collected by trypsinization, neutralized in serum-containing DMEM, and lysed in a 1:1 mixture of methanol and 1 N perchloric acid (MeOH-PCA) on ice for 10 min. Lysates were centrifuged at 10,000 × g for 10 min at 4 °C, and PPIX fluorescence was measured in black 96-well plates under identical spectrophotometric settings. In all cases, PPIX concentrations were determined using a standard curve prepared with PPIX (Sigma-Aldrich) and normalized to tissue weight or total protein content in each sample as measured by BCA.

### Statistics

All quantitative experiments were performed using at least three independent biological replicates. Data are presented as mean ± standard error of the mean (SEM). Statistical analyses were conducted using GraphPad Prism v9 (GraphPad Software). For pairwise comparisons, Student’s t-tests were used. For multiple group comparisons, one-way ANOVA followed by Tukey’s post hoc test was applied. A p-value of <0.05 was considered statistically significant. Statistical outcomes are provided in the figure legends. No data points were excluded from the analyses.

## Data availability statement

All data supporting the findings of this study are available within the article and its Supplementary Materials.

## Acknowledgements

We are thankful for fellowship support from the Vietnam Education Foundation to Lan N. Tu and for partial research support from the Dextra Undergraduate Research Fund at Cornell University to Carmen Smith. We also thank Meghan Showalter and Oliver Fiehn at UC Davis for lipidomic services. The authors gratefully acknowledge the contributions of all current and former members of the Selvaraj Laboratory of Integrative Physiology, whose efforts and insights have shaped the direction of this work. We further thank Dr. Yves Boisclair and his laboratory at Cornell for valuable discussions that helped refine the conceptual framework of this study and enabled some of the discoveries presented here.

## Supplementary Information

**Table S1.**
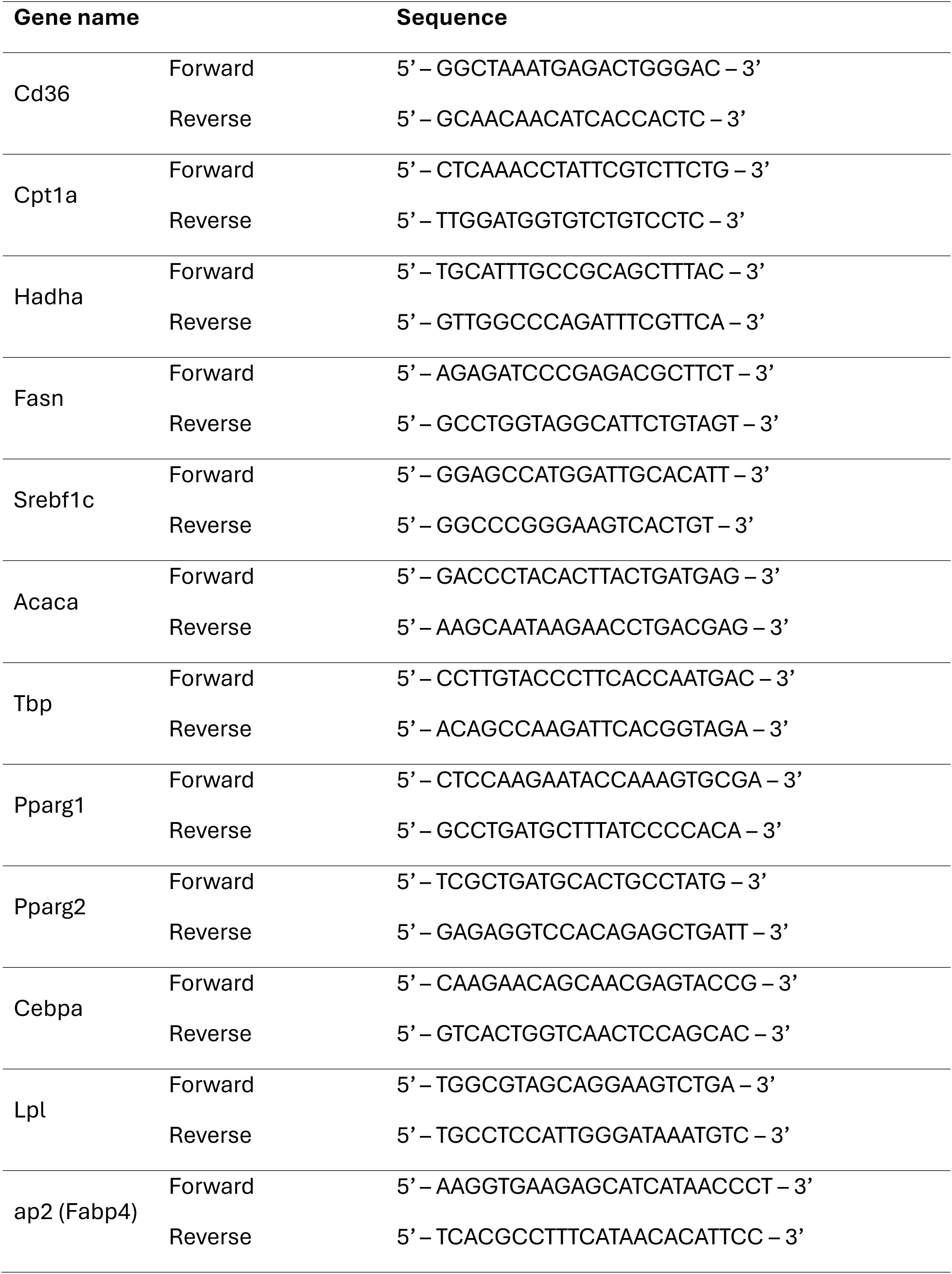
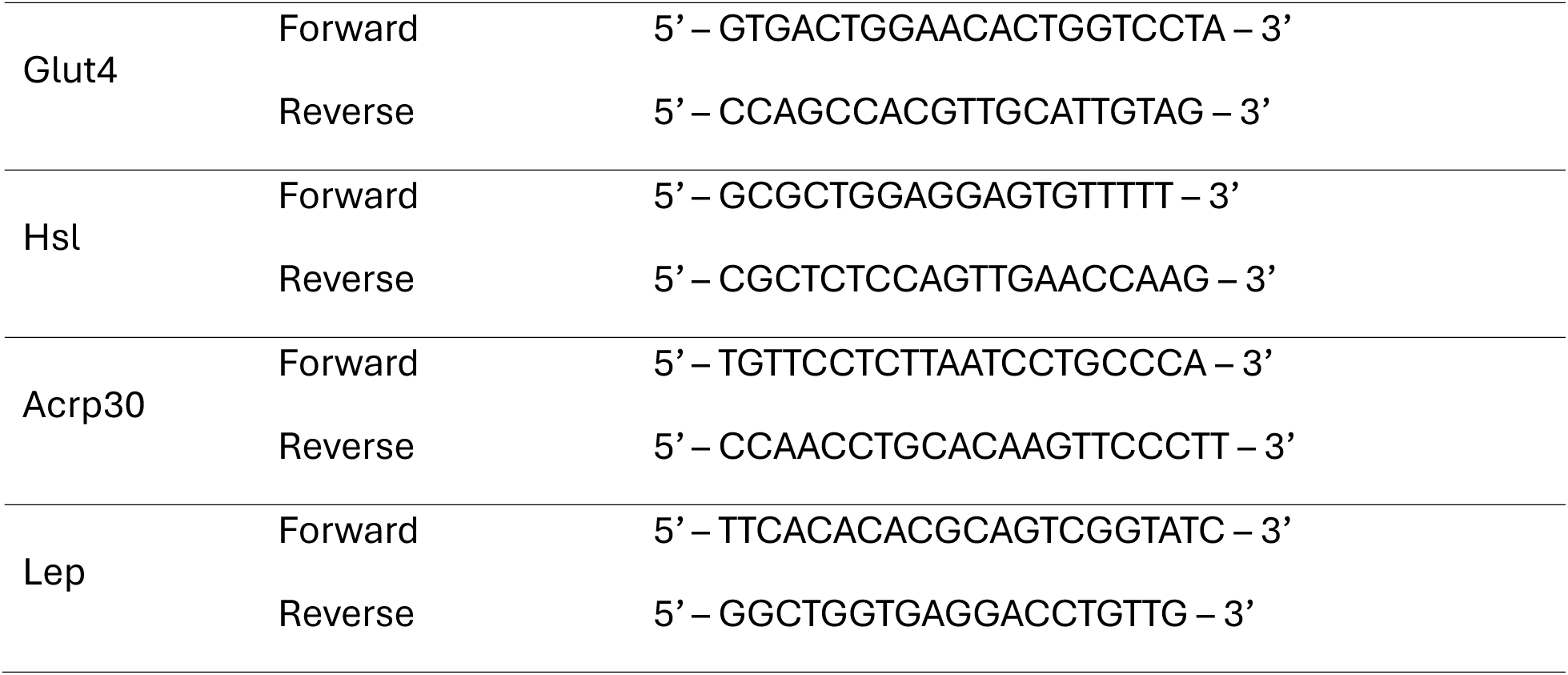
Primers used for gene expression.

